# Repeat Opioid Use Modulates Microglia Activity and Amyloid Beta Clearance in a Mouse Model of Alzheimer’s Disease

**DOI:** 10.1101/2025.11.14.688256

**Authors:** Joseph V Gogola, Sung Won Stephanie Wee, Alfredo J Garcia

**Affiliations:** Department of Medicine, The University of Chicago, Chicago IL USA; Department of Neurobiology, The University of Chicago, Chicago IL USA; Biosciences Division, Argonne National Laboratory, Lemont IL USA

**Keywords:** Opioids, Fentanyl, Chronic Drug Exposure, Neuroinflammation, Alzheimer’s Disease, Microglia

## Abstract

In addition to driving dependency and overdose, illicit use of opioids, such as fentanyl, is linked to the risk for cognitive decline and dementia. Growing evidence also indicates that opioid use is associated with pathological features, paralleled early in Alzheimer’s disease (AD), which raises the possibility of the involvement of mechanistic interactions between opioid use and AD progression. Here, we investigate how chronic fentanyl use (i.e., 20 days) influences the neuroimmune state, microglial activity, and amyloid burden in wildtype and APPPS1-21 mice, a transgenic model of AD. In wild-type mice, fentanyl use promoted a pro-inflammatory state without increasing the incidence of disease-associated microglia. In APPPS1-21 mice, chronic fentanyl use led to a shift favoring an anti-inflammatory state, which was associated with increased microglia clustering and activation at Aβ plaques, increased Aβ internalization in plaque-associated activated microglia, decreased soluble Aβ, and decreased plaque burden. Our findings indicate that chronic fentanyl use fundamentally changes the trajectory of neuroimmune activity and features characteristic of early AD by enabling microglia to enhance Aβ clearance. The interactions demonstrate how substance use can reshape the neuroimmune landscape in neurodegenerative disease, emphasizing the importance of tailored treatment strategies.

## Introduction

The ongoing opioid epidemic poses a significant threat to public health, affecting millions with opioid dependence across the United States and globally. Although opioids continue to play a crucial role in managing acute pain and providing palliative care, prolonged usage has demonstrated concerning neurological consequences that extend well beyond the established risks of dependency and overdose-related complications. Large-scale population analyses indicate that sustained opioid treatment increases the likelihood of developing dementia across all subtypes, with vascular dementia showing the most pronounced correlation (Dublin et al., 2015; Gao et al., 2024; Billioti de Gage et al., 2012; Lin et al., 2025). Brain imaging studies have also documented structural changes linked to opioid use, including decreased volume in the hippocampus and white matter regions, indicating potential direct effects on brain deterioration in areas vital for cognitive function and memory formation (Warner et al., 2024; Upadhyay et al., 2010; Gao et al., 2024; Lin et al., 2025). While these features parallel the initial stages of Alzheimer’s disease (AD) development (Gao et al., 2024; Younger et al., 2009; Liang et al., 2016), these correlations between opioids and cognitive decline raise the pressing question: How does chronic fentanyl use influence the development or advancement of AD?

Here, we examine how chronic opioid use affects amyloid plaque burden, microglial activity, and cytokine signaling in a mouse model of AD. Opioids directly modulate microglial function through mu-opioid receptors (MOR), with chronic exposure inducing microglial reactivity that affects a variety of inflammatory and behavioral features (Green et al., 2022). Given that microglial activation represents a central mechanism in AD pathogenesis – where these cells regulate amyloid-β plaque dynamics through inflammatory signaling cascades – we hypothesized that chronic opioid administration would interfere with the microglial response in AD, thereby increasing Aβ plaque burden and associated disease markers. Surprisingly, we found the opposite. Repeated opioid use (ROU) led to decreased plaque burden, increased microglial clustering and activation in the plaque vicinity in APPPS1-21 mice, a model for AD. These changes were corresponded with a decrease in soluble but not insoluble Aβ and evidence of increased Aβ internalization by plaque-associated microglia. Critically, these putatively beneficial effects of repeated opioid use coincided with a shift toward anti-inflammatory signaling, contrasting with the pro-inflammatory response observed in wild-type mice receiving the same treatment. These data suggest that the pre-existing pathological state of the AD brain fundamentally alters how it responds to opioid exposure, potentially converting what would typically be a detrimental inflammatory response into a beneficial one for augmenting Aβ clearance via increased microglial phagocytosis of amyloid plaques. These previously undescribed phenomena demonstrate the impact of chronic opioid use on amyloid burden and the inflammatory state relevant to AD pathology and underscore the necessity to consider a history of opioid use.

## Results

### Fentanyl Treatment Promotes Pro-Inflammatory Response in Wild-Type Mice Without Altering Microglial Coverage

To establish the effects of chronic opioid exposure, we first examined the response in wild-type (“WT”) mice. We administered fentanyl (“ROU”) or saline (“Sal”) daily to male wild-type mice, starting at postnatal day (P) 130 and ending at P150. Microglia coverage did not differ between treatment groups in either cortex (**Figure 1A-C**; WT-Sal = 3.847±0.44, WT-ROU = 3.67±0.406, *p* = 0.9397) or hippocampus (WT-Sal = 2.651±0.41, WT-ROU = 2.37±0.161, *p* = 0.7918), suggesting no fentanyl-induced effect on gliogenesis. Fentanyl treatment led to an elevation in the pro-inflammatory marker, TNF-α (**Figure 1D-E**; WT-Sal = 2.018±0.33; WT-ROU = 7.87±0.572, *p* = 0.00002), without changing the anti-inflammatory marker IL4 (**Figure 1F-G**; WT-Sal = 1.864±0.276, WT-ROU = 2.066±0.285, *p* = 0.6236). These results indicate that chronic fentanyl exposure shifts the neuroinflammatory environment toward a more pro-inflammatory state in wild-type mice without affecting basal microglial distribution.

**Figure 1.**
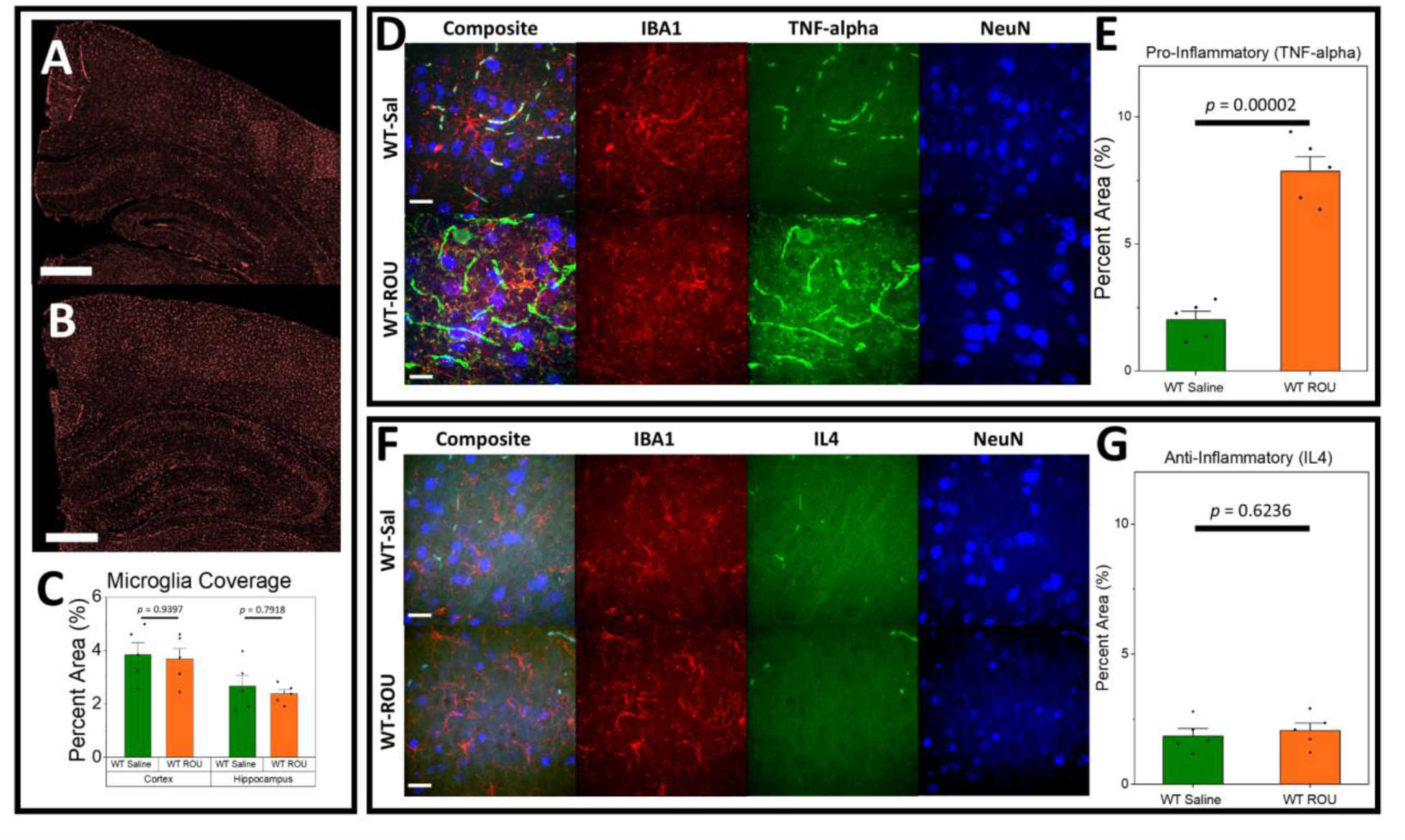
ROU Leads to Proinflammatory Response in Wild-Type Mice. **(A-C)** There are no broad differences in microglia coverage between Saline-(**A**) or ROU-treated (**B**) wild-type mice. **(D-E)** Pro-inflammatory marker levels are significantly increased with ROU treatment. **(F-G)** Anti-inflammatory levels are unchanged with ROU treatment. Data shown as mean ± SEM, N=5 per group. Scale bar = 500 μm (A, B) or 20 μm (D, F).

### Fentanyl Treatment Shifts APPPS1-21 Mice Toward Anti-inflammatory Profile

We next examined the cytokine response to chronic fentanyl treatment in APPPS1-21 mice (“AD”) (**Figure 2A**). In contrast to WT mice, fentanyl treatment in APPPS1-21 mice showed no change in TNF-α (**Figure 2B**; AD-Sal = 14.601±2.952, AD-ROU = 12.395±3.167, *p* = 0.1344), though at a higher baseline than wild-type mice. Surprisingly, ROU produced a significant increase in IL-4 in AD mice (AD-Sal = 13.862±1.886, AD-ROU = 26.427±4.224, *p* = 0.000178). This result was particularly visually pronounced in the regions immediately surrounding amyloid plaques (**Figure 2A**). Moreover, fentanyl treatment had no significant effect on microglia coverage broadly in either cortex (**Figure 2C**; AD-Sal = 10.229±1.047, AD-ROU = 9.959±0.558, *p* = 0.9951) or hippocampus (AD-Sal = 8.148±1.008, AD-ROU = 7.827±0.510, *p* = 0.9918).These results demonstrate that the pre-existing pathological state of the APPPS1-21 brain fundamentally alters both its baseline inflammatory state, as well as how it responds to opioid exposure, shifting toward a more anti-inflammatory state rather than the pro-inflammatory response observed in wild-type mice.

**Figure 2.**
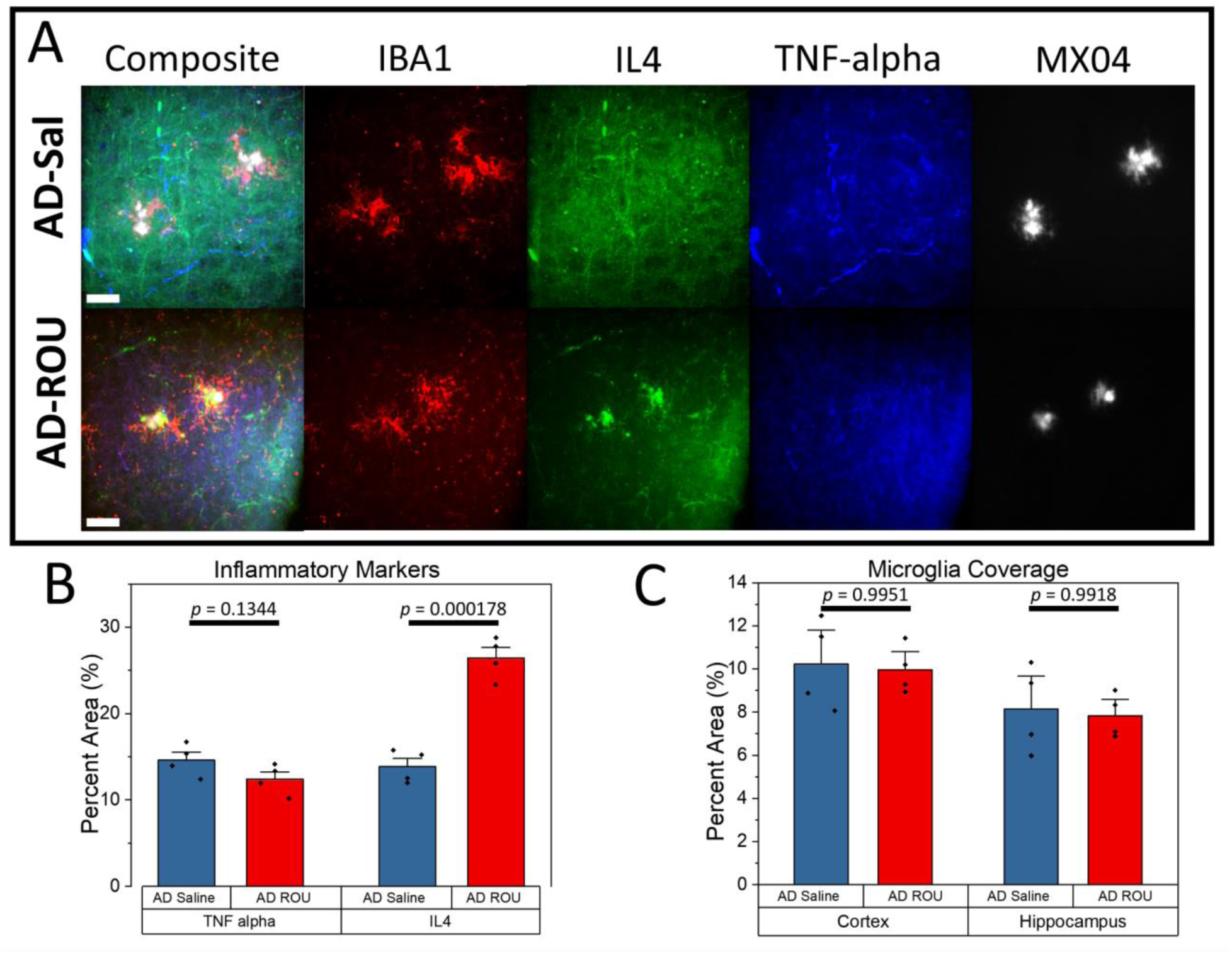
Repeated opioid use increases Anti-inflammatory state without altering Pro-inflammatory state or gliogenesis in AD mice. (**A**) Representative immunofluorescence images of microglia (IBA1), IL-4, TNF-α, and amyloid plaques (MX04) in cortex of saline-treated (AD-Sal) and repeated fentanyl-treated (AD-ROU) APPPS1-21 mice. (**B**) Quantification of inflammatory marker coverage (% area). AD-ROU mice showed significantly increased IL-4 coverage compared to AD-Sal, particularly around amyloid plaques, while TNF-α coverage was unchanged. (**C**) Quantification of microglia coverage shows no treatment effect in either cortex or hippocampus. Data shown as mean ± SEM, N=4 per group. Scale bars = 25 μm.

### Microglial Recruitment and Activation at Plaques is Enhanced Following Fentanyl Treatment in APPPS1-21 Mice

Given the differential inflammatory responses between WT and APPPS1-21 mice, we examined whether fentanyl treatment also affected microglial behavior around amyloid plaques. One of the best documented cellular features of Aβ plaques is microglial clustering around these local lesions. Using standard immunofluorescent labeling of Iba1-positive microglia, we found fentanyl-induced changes in microglial coverage at plaques in APPPS1-21 mice (**Figure 3A**). Fentanyl-treated mice showed significantly greater Iba1+ coverage at plaques compared to saline controls in both cortex (**Figure 3B**; AD-Sal = 38.67±1.97, AD-ROU = 66.73±1.59, *p* = 1.6E-6) and hippocampus (AD-Sal = 22.16±1.41, AD-ROU = 61.33±1.13, *p* = 3.7E-8). To assess microglial activation status, we also co-labeled with Clec7a as a marker of disease-associated microglia. Clec7a signal was almost exclusively restricted to the immediate peri-plaque locations in both treatment groups of APPPS1-21 mice (Figure 3A), and ROU treatment led to no Clec7a+ microglia expression in WT mice (data not shown). Fentanyl-treated AD mice showed significantly greater Clec7a+ coverage at plaques compared to saline controls in both cortex (**Figure 3C**; AD-Sal = 35.79±2.80, AD-ROU = 63.75±1.97, *p* = 2.39E-5) and hippocampus (AD-Sal = 15.59±1.48, AD-ROU = 56.13±1.38, *p* = 3.8E-7). Surprisingly, measures of plaque coverage for both total and activated microglia were similar after fentanyl treatment, despite differing baseline coverage levels between cortex and hippocampus in the saline control group (Iba1: *p* = 0.014; Clec7a: *p* = 0.018). Thus, chronic fentanyl exposure promotes both microglial recruitment and microglial activation at plaques in the AD brain.

**Figure 3.**
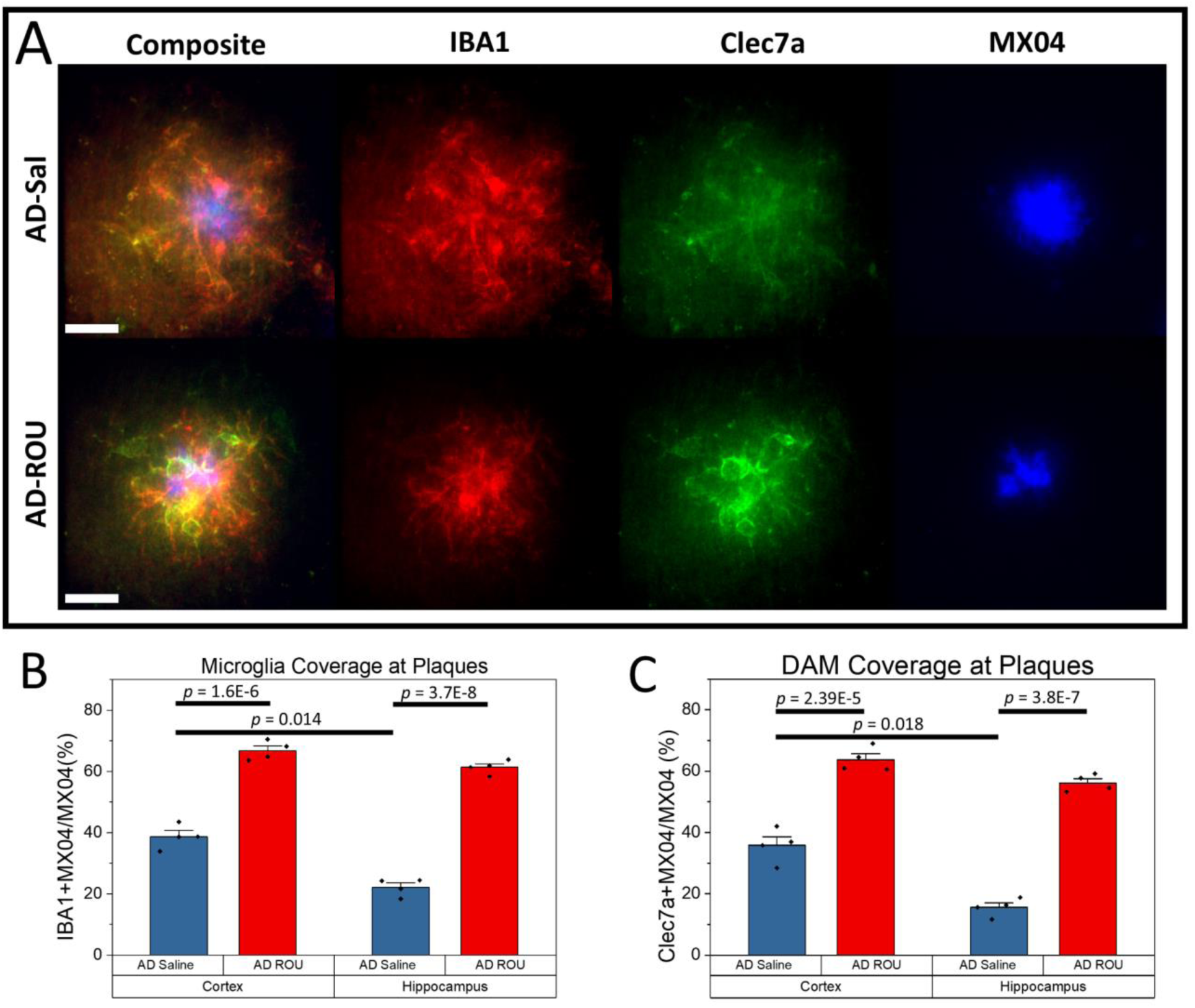
Repeated opioid use increases microglia recruitment and activation at amyloid plaques. (**A**) Representative confocal images of cortical amyloid plaques in saline-treated controls or ROU-treated AD mice. Red, total microglia; green, activated microglia; blue, amyloid plaques. (**B-C**) Quantification of plaque coverage by (B) total microglia (IBA1+ area overlapping MX04+) and (C) activated microglia(Clec7a+ area overlapping MX04+) in cortex and hippocampus. AD-ROU mice exhibited significantly increased total and activated microglia at plaques compared to AD-Sal controls. Data shown as mean ± SEM, N=4 per group. Scale bars = 20 μm.

### Enhanced Microglial Phagocytosis and Reduced Soluble Amyloid-β Following Fentanyl Treatment

Since chronic fentanyl administration increased activated microglia at plaques, we next investigated whether this translated to enhanced amyloid clearance. Analysis of high-magnification images from the immunofluorescent samples revealed internalized Aβ within plaque-associated activated microglia (**Figure 4 A-B**; gold arrows). Quantification of intracellular Aβ signal within Clec7a+ microglia at plaque sites showed significantly higher internalized Aβ in the fentanyl treatment group compared to saline controls, in both cortex (**Figure 4C**; AD-Sal = 11.22±0.60, AD-ROU = 19.11±0.99, *p* = 0.0016) and hippocampus (AD-Sal = 4.10±0.35, AD-ROU = 17.04±0.76, *p* = 4.6E-5). We saw no difference in internalized Aβ signal between brain regions in the ROU group, despite differing baselines in the saline control (*p* = 0.0002). These results suggest that fentanyl treatment is inducing peri-plaque activated microglia to phagocytose Aβ.

**Figure 4.**
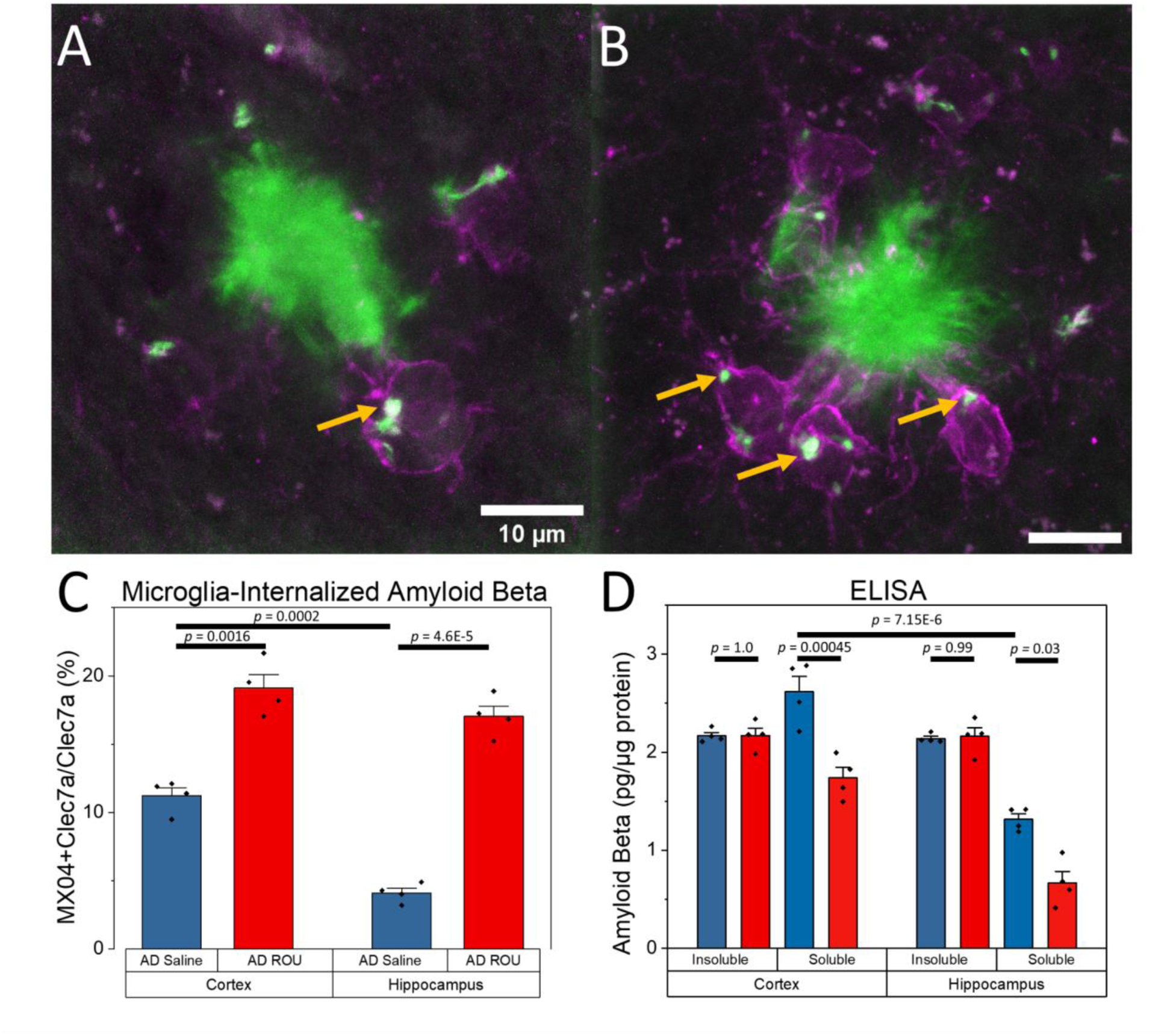
Repeated opioid use enhances microglial Aβ clearance in AD mice. (**A–B**) Representative images of Aβ (MX04+; green) and activated microglia (Clec7a+; magenta) in cortex of AD mice treated with saline (A) or fentanyl (B). Microglia in AD-ROU show increased internalization of Aβ (gold arrows). (**C**) Internalized Aβ as a proportion of total Clec7a+ microglia at the plaque, in cortex and hippocampus. AD-ROU mice show increased Aβ uptake compared to AD-Sal controls. (**D**) Insoluble and soluble Aβ levels in cortex and hippocampus. AD-ROU treatment reduced soluble Aβ in both regions, with no effect on insoluble amyloid beta. Data shown as mean ± SEM, N=4 per group. Scale bar = 10 μm.

To further assess potential amyloid clearance mechanisms, we performed Enzyme-Linked Immunosorbent Assays (ELISA) for soluble and insoluble Aβ species. ELISA analysis revealed distinct effects of fentanyl treatment on soluble versus insoluble Aβ pools (**Figure 4D**). Soluble Aβ levels were significantly reduced in fentanyl-treated AD mice compared to saline controls in both cortex (AD-Sal = 2.62±0.16 pg/μg protein, AD-ROU = 1.74±0.11, *p* = 0.00045) and hippocampus (AD-Sal = 1.32±0.06, AD-ROU = 0.67±0.12, *p* = 0.03). In contrast, insoluble amyloid-β concentrations remained unchanged between treatment groups in both cortex (AD-Sal = 2.16±0.03, AD-ROU = 2.17±0.07, *p* = 1.0) and hippocampus (AD-Sal = 2.14±0.02, AD-ROU = 2.16±0.09, *p* = 0.99). Of note, we found no ROU-induced changes in soluble or insoluble Aβ in the WT mice, where measures for most samples were at or below detection threshold (data not shown). These findings indicate that chronic fentanyl exposure specifically affects the soluble amyloid pool in the AD brain, consistent with enhanced clearance mechanisms rather than reduced amyloid generation broadly. The enhanced internalization of Aβ by activated microglia (**Figure 4A-B**) is consistent with the decrease in soluble Aβ levels, suggesting peri-plaque activated microglia are actively phagocytosing Aβ from plaques to restrain ongoing aggregation.

### Chronic Fentanyl Treatment Reduces Amyloid Plaque Burden in APPPS1-21 Mice

Finally, we examined the ultimate outcome of enhanced microglial clearance on overall amyloid pathology. Methoxy-X04 staining of Aβ plaques (**Figure 5A-B**) revealed that fentanyl-treated APPPS1-21 mice exhibited significantly reduced area-normalized plaque number compared to saline-treated controls in both cortex (**Figure 5C**; AD-Sal = 143.19±2.90 plaques/cm^2^, AD-ROU = 105.01±2.46, *p* = 0.00029) and hippocampus (AD-Sal = 62.64±5.33, AD-ROU = 22.98±1.49, *p* = 0.00023). Further analysis confirmed that plaque coverage, or the relative proportion of each brain region positive for amyloid plaques, was also decreased in the ROU group relative to saline controls for both cortex (**Figure 5D**; AD-Sal = 0.495±0.039, AD-ROU = 0.321±0.029, *p* = 0.016) and hippocampus (AD-Sal = 0.307±0.024, AD-ROU = 0.114±0.021, *p* = 0.0098). This pattern was consistent across both brain regions examined, despite differing plaque burden and coverage between the two regions in control mice (Plaque Burden: *p* = 3.17E-6; Plaque Coverage: *p* = 0.00085). Of note, we found no incidence of Aβ plaques in WT mice, in either treatment group (data not shown). These results demonstrate that the enhanced microglial activation and clearance mechanisms induced by chronic fentanyl exposure ultimately translate to reduced amyloid plaque burden in the AD brain, suggesting a protective effect against amyloid pathogenesis.

**Figure 5.**
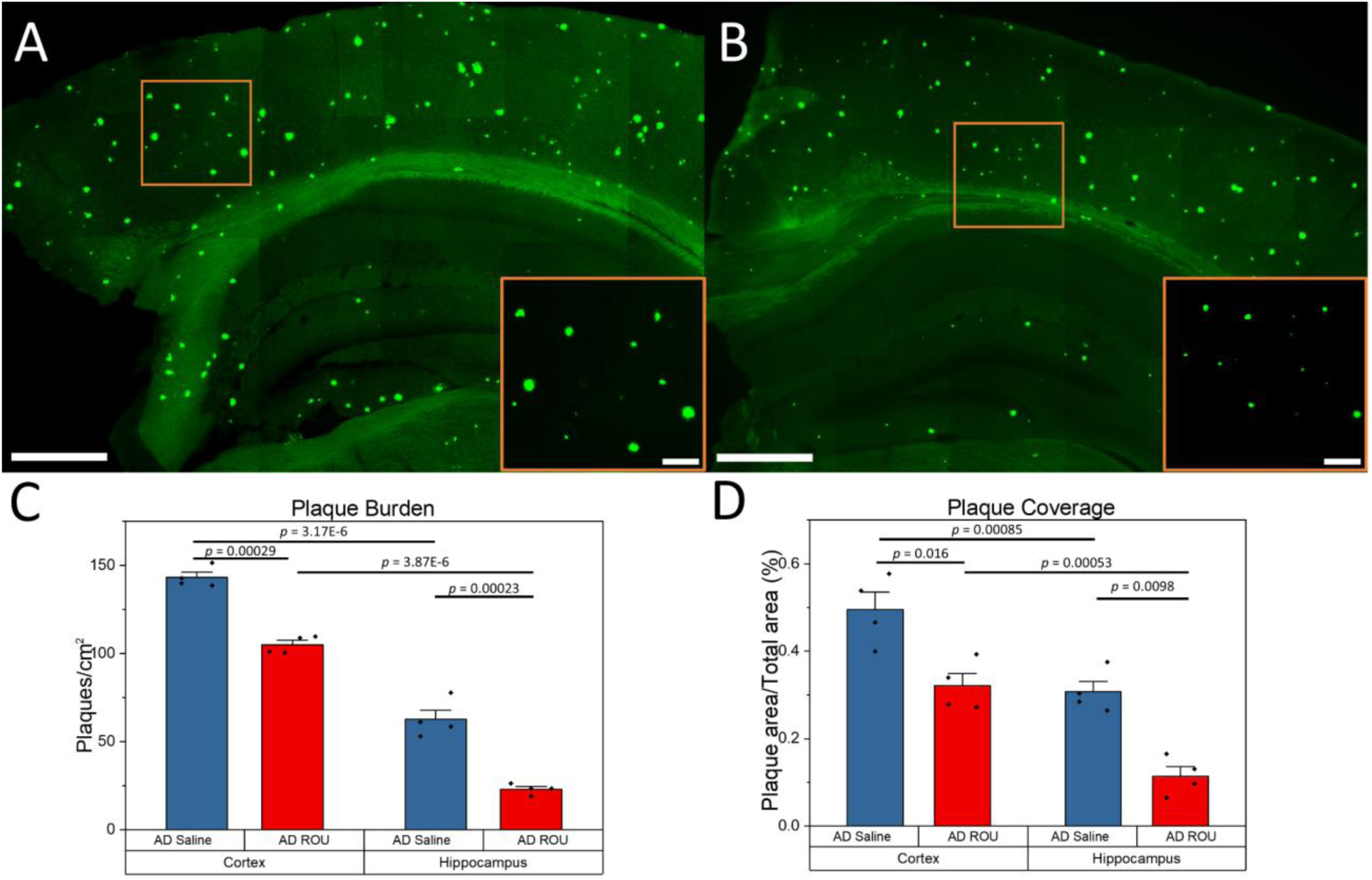
ROU treatment reduces amyloid plaque accumulation in AD mouse brain. (**A–B**) Coronal brain sections from AD mice labeled with Methoxy-X04 (green). Compared to saline-treated controls (A), ROU-treated mice (B) show fewer and smaller plaques in both cortex and hippocampus. (**C**) Quantification of plaque burden, expressed as the number of plaques per cm². ROU treatment significantly lowered plaque burden in both cortex and hippocampus compared to saline controls. (**D**) Quantification of plaque coverage, expressed as the percentage of tissue area occupied by plaques. ROU treatment reduced the proportion of cortex and hippocampus covered by plaques. Data shown as mean ± SEM, N=4 per group. Scale bars = 500 μm (overview) or 100 μm (gold-inset).

## Discussion

Beyond the immediate risks of addiction and overdose, emerging evidence links chronic opioid exposure to accelerated cognitive decline (Bhatia et al., 2023; Warner et al., 2022) and increased risk for dementia (Gao et al., 2024; Pourhadi et al., 2024), creating a dual public health crisis at the intersection of substance use and neurodegenerative disease. The illicit fentanyl epidemic is responsible for over 70% of overdose deaths in the U.S., with life expectancy dropping in affected populations. AD affects over 6 million people in the U.S. alone, with aging populations are expected to double this burden by mid-century. Both are independently devastating, and, increasingly intersecting – epidemiologically, mechanistically, and socially – because older adults, the demographic most vulnerable to AD, represent a growing proportion of opioid users (Shoff et al., 2021; Dufort and Samaan, 2021; Kuo et al., 2025).

Against this backdrop of well-documented harm, our findings reveal a striking and unexpected pattern: chronic fentanyl use enhances microglial activation and improves amyloid beta clearance in a mouse model of Alzheimer’s disease. This pattern is characterized by an enhanced anti-inflammatory response, despite promoting pro-inflammatory responses in wild-type mice, suggesting that the effect of fentanyl on neuroimmune status is influenced by the presence of pathohistological biomarkers of AD. This challenges the conventional assumptions about opioid-disease interactions, e.g., that opioids are neurotoxic and exacerbate disease pathology. Furthermore, this emphasizes the significance for considering the interplay between the effects of drug use on underlying neurodegenerative conditions.

Microglia are key mediators of opioid-driven neuroinflammation that can undermine analgesic efficacy. Pharmacological inhibition or depletion of microglia modifies opioid-induced tolerance and hyperalgesia. For example, minocycline, a drug commonly used to suppress microglial activation, significantly attenuates morphine tolerance by suppressing p38 MAPK phosphorylation and pro-inflammatory cytokine production in spinal microglia (Cui et al., 2008; Mika et al., 2009). Microglial depletion or inhibition has been reported to attenuate morphine-induced hyperalgesia and markers of central sensitization in some pain models, and to alter tolerance development when pre-administered prior to chronic opioid exposure (Ferrini et al., 2013; Grace et al., 2016; Reiss et al., 2020; Hayashi et al., 2016). While these results implicate microglia as mediators of opioid-driven neuroinflammation and maladaptive responses that oppose analgesia in otherwise healthy nervous systems, they also support the view that fentanyl may act to affect the microglial response to neurodegenerative conditions.

While the basis by which fentanyl mediates changes to microglial activity remains to be further resolved in APPPS1-21 mice, opioids have been shown to interact with toll-like receptor 4 (TLR4), a key activator of the innate immune response that is primarily expressed in microglia (Lehnardt et al., 2003). Fentanyl and other clinically relevant opioids bind to TLR4 by docking to the lipopolysaccharide (LPS)-binding pocket of MD-2 (Wang et al., 2012; Hutchinson et al., 2009), triggering MyD88-dependent activation of NF-κB and subsequent transcription of pro-inflammatory cytokines (TNF-α, IL-1β, IL-6) (Xie et al., 2017; Grace et al., 2016). Pharmacological blockade of TLR4 signaling or genetic ablation of the receptor potentiate opioid analgesia, attenuate the development of tolerance and hyperalgesia, and reduce withdrawal behaviors (Eidson & Murphy, 2013; Ferrini et al., 2013; Hutchinson et al., 2010; Reiss et al., 2020; but see Liu et al., 2022), supporting the potential that interaction between fentanyl and TLR4 may be critical to the neuroimmune response observed here.

Amyloid plaques themselves function as endogenous TLR4 agonists, creating a fundamentally different baseline inflammatory state in the AD brain that transforms how chronic fentanyl exposure is interpreted by the microglial compartment. Fibrillar Aβ binds to and activates TLR4, initiating intracellular signaling cascades including Src-Vav-Rac activation, p38 MAPK phosphorylation, reactive oxygen species production, and phagocytosis (Reed-Geaghan et al., 2009; Yu et al., 2012). This plaque-driven TLR4 priming creates a chronic inflammatory environment analogous to that induced by LPS, the prototypical TLR4 agonist. Critically, the presence of Aβ increases TLR4 expression (Calvo-Rodríguez et al., 2017), amplifying potential Aβ interactions, inflammatory outcomes, and phagocytic capacity.

Microglia lacking TLR4 fail to bind fibrillar Aβ and cannot initiate phagocytosis (Reed-Geaghan et al., 2009), as do those lacking key molecular co-factors such as Lyn kinase (Islam et al., 2025). Microglial TLR4 loss-of-function mutations also increase Aβ deposits and exacerbate cognitive deficits (Song et al., 2011). Moreover, microglial depletion studies demonstrate that in the absence of microglia, Aβ plaque compaction is compromised, resulting in more diffuse plaque morphologies and increased neuritic dystrophy, effects which can be reversed with microglial repopulation (Casali et al., 2020). Thus, microglia substantially contribute to Aβ pathogenesis, with TLR4 situated as both a key mediator for these processes and a lodestone that may be exploited by fentanyl or other opioids.

We propose that, in a plaque-primed state, the interaction between fentanyl and microglia may shift TLR4 signaling into a state favoring phagocytic clearance over persistent inflammation. This would be reminiscent of endotoxin tolerance, where repeated TLR4 activation shifts responses from pro-inflammatory to regulatory or anti-inflammatory phenotypes (Bohannon et al., 2014; O’Carroll et al., 2014; Vergadi et al., 2018). Indeed, the microglial response to LPS is muted in aged AD mice, where activation corresponds to reduced Aβ burden (Go et al., 2016). Given that plaque-driven TLR4 activation increases TLR4 receptor expression, fentanyl may act as a secondary signal that amplifies clearance mechanisms, rather than initiating *de novo* inflammation. The chronically activated baseline may also modify MOR signaling dynamics, potentially shifting receptor coupling toward anti-inflammatory pathways (Miao et al., 2023) or altering MOR–TLR4 crosstalk (Gessi et al., 2016; Franchi et al., 2012). Microglial-specific manipulations – for example, conditional perturbations of microglial MOR or TLR4 – would help clarify whether the putatively beneficial effects observed here require microglial TLR4 signaling, MOR-mediated pathways, or context-specific interactions between these systems.

Microglia exhibit remarkable plasticity and are central to Aβ plaque dynamics (Prinz et al., 2019). Disease-associated microglia (DAM) represent a specialized subset often localizing around Aβ plaques (Keren-Schaul et al., 2017). DAMs express characteristic markers including Clec7a (Dectin-1), which is associated with enhanced phagocytic capacity and Aβ clearance, making it a key indicator of microglial activation state around plaques (Zhao et al., 2023; Litvinchuk et al., 2018). The clustering of microglia around plaques and their expression of DAM markers like Clec7a correlate with plaque internalization and clearance, representing a protective mechanism against plaque accumulation (Hong et al., 2016; Zhao et al., 2023). Enhanced microglial clustering and Clec7a expression following fentanyl treatment suggest that opioids may activate endogenous protective mechanisms rather than simply suppress inflammation.

Our demonstration of increased Aβ internalization in plaque-associated Clec7a+ microglia provides direct evidence for such enhanced clearance. This, coupled with reduced soluble but not insoluble Aβ levels, supports the notion that chronic fentanyl enhances removal of Aβ plaques rather than reducing Aβ production (Hong et al., 2016; Griciuc et al., 2019). Thus, opioid exposure may be stimulating DAM signaling pathways to promote beneficial microglial activation for post-depositional clearance rather than prevention of Aβ aggregation or a decreased production of Aβ species more broadly.

While we modeled how fentanyl use (as occurs often in opioid abuse and opioid use disorder) affects Aβ burden, and found that enhanced Aβ clearance may result, these findings might support the potential therapeutic value of opioids at clinically relevant doses in reducing Aβ accumulation. While improved Aβ clearance represents a promising outcome, the functional consequences for cognitive performance and behavioral symptoms remain unexplored. Future studies must determine whether enhanced plaque clearance translates to meaningful improvements in learning, memory, and other cognitive domains affected in AD, as well as other biomarkers of disease progression.

The observed effects in both hippocampus and cortex – two regions critically affected in AD – suggest that opioid-induced enhancement of microglial function is not limited to specific brain areas but may represent a more generalized protective response in the diseased brain (Benzinger et al., 2013; Furcila et al., 2019). However, plaques emerge at different times in these areas: neocortex at ∼P42 and hippocampus between P84–P123 (Radde et al., 2006). Thus, in relative terms, plaques in cortex might be considered older than plaques in hippocampus at our collection timepoint, and relative contributions of microglia in earlier vs. later pathology are well documented (d’Errico et al., 2021; Sun et al., 2023; Baligács et al., 2024). Thus, our finding that microglial coverage, activation, and internalization after fentanyl treatment are similar across cortex and hippocampus, despite baseline differences, suggests that the capacity for alterations in microglia activation and functional state might be similar despite these relative differences in plaque age. These regional differences raise important questions about optimal targeting strategies, as well as dose– and age-dependent effects. The shift toward smaller plaque sizes likely reflects enhanced clearance of smaller plaques, and/or compaction of existing plaques through increased microglial containment, both of which could represent beneficial outcomes but through different mechanisms. Understanding these regional variations will be crucial for any potential therapeutic applications.

Our results fundamentally challenge the traditional view that drug use is universally detrimental in neurodegenerative diseases (Volkow & Koob, 2015; Billioti de Gage et al., 2012; Rehm et al., 2019; Dublin et al., 2015; Gao et al., 2024). The beneficial effects we observed suggest that pathological brain states may alter drug responses in ways that convert typically harmful exposures into potentially therapeutic ones (Kato et al., 1977; Spedding et al., 2005). This paradigm shift has broad implications beyond opioids and AD, suggesting that other drug-disease interactions may need reevaluation in light of baseline pathological states.

The implications of these findings are complex and require careful consideration. While our results suggest potential therapeutic benefits, they do not constitute a recommendation for opioid use in AD patients, given the substantial risks of addiction, tolerance, and other adverse effects (Volkow & Koob, 2015). Importantly, the dose given here (0.35 mg/kg) would be characterized as a high or abusive dose, not a therapeutic dose typically given in a clinical setting, which tend to be orders of magnitude smaller (Sobieraj et al., 2019; Fujii et al., 2019). Dose-response relationships need careful characterization to identify potential therapeutic windows. The beneficial effects we observed may only occur within a specific dose range, with higher doses potentially causing toxicity or impairing microglial function, and lower doses failing to reveal the beneficial effects on plaque clearance.

Additionally, our study examined only acute effects of repeated opioid exposure, and the long-term consequences remain unknown. It is critical to determine whether the beneficial effects on Aβ clearance persist with continued treatment, reverse upon opioid cessation, or represent transient changes that could inform optimal dosing strategies. Finally, the translation from mouse models to human AD patients requires consideration of species differences in opioid metabolism, receptor expression patterns, and disease progression timelines (Sasaguri et al., 2017; Kato, 1977; Do Carmo and Cuello, 2013). Future studies using more clinically relevant dosages – such as those achieved with partial agonists like buprenorphine, which are already used safely in substance use disorder treatment (Shulman et al., 2019) – would be essential towards realizing the possibility of developing opioid-inspired therapeutics that could capture the beneficial microglial activation effects while avoiding the addiction liability and other systemic complications associated with traditional opioids and their abuse.

In conclusion, our study reveals an unexpected beneficial interaction between repeated opioid use and AD pathology, mediated through enhanced microglial activation and Aβ clearance. While these findings do not support clinical opioid use in AD patients, they provide novel insights into context-dependent drug responses and highlight the therapeutic potential of targeting microglial function through carefully designed pharmacological interventions. While future research must resolve the mechanistic basis of our observations, our results emphasize the critical importance of considering baseline disease states when evaluating any therapeutic intervention (Kato, 1977; Ashley, 2016).

These insights could extend beyond AD to other neurodegenerative conditions characterized by microglial dysfunction and protein aggregation (Song & Colonna, 2018; Subhramanyam et al., 2019; Hou et al., 2021). Thus, the differential responses we observed here support the need to account for the interaction of substance abuse disorder with other underlying pathological conditions, such as AD.

## Methods

### Mouse line

Transgenic mice known as APPPS1-21, which contain human transgene for both amyloid precursor protein (APP) with the Swedish mutation and presenilin1 (PSEN1) with the L166P mutation, were used in this study (JAX stock #037891). All animals used in this study were maintained on a hybrid background generated by crossing female C57BL/6J with male APPPS1-21. Male APPPS1-21 mice, aged P130 (n=8), along with their wild-type littermates (n=10) were randomly assigned to treatment groups. All mice were housed in groups under the same 12:12h light-dark cycle with *ad libitum* access to food and water. All procedures were performed with the approval of the Institute of Animal Care and Use Committee at The University of Chicago.

### Repeated Opioid Use and Fentanyl Administration

The repeated opioid use (ROU) protocol consisted of daily IP injections of fentanyl (0.35mg/kg) for 20 consecutive days in both wild-type and APPPS1-21 mice. Control groups received an equivalent volume of saline (IP injection) in place of fentanyl. Brain tissues were harvested approximately 24 hours after the final injection. Briefly, animals were anaesthetized with isofluorane, then perfused first with 0.01 M phosphate buffered saline (PBS) followed by 4% paraformaldehyde in PBS. Brains were post-fixed overnight at 4°C in fixative solution, then cryoprotected in 30% sucrose, sectioned at 40µm thick on a Cryostat, and stored in antifreeze at –20°C until further processing.

### Immunohistochemistry

Immunohistochemical staining was performed to assess Aβ deposition, microglial activation, and expression of cytokines in mouse brain tissue. Brain sections were washed with PBS and then blocked using 10% normal donkey serum (JacksonImmuno Research, 017-000-121) in PBS supplemented with 0.1% Triton X-100 (PBS-T, Sigma, T8787-50mL). Sections were incubated overnight at 4°C with primary antibodies diluted in the blocking buffer. Primary antibodies used in this study are listed in **Table 1**. After washing in PBS-T and PBS to remove unbound antibodies, sections were incubated with the matching secondary antibodies (listed in **Table 2**) in blocking buffer, including methoxy-X04. Sections were further rinsed in PBS-T and PBS and lastly stained with TOPRO (DF). Slides were cover slipped with Fluoromount-G^TM^ Mounting Medium (Invitrogen, Cat# 00-4958-02) for fluorescence imaging and stored at 4°C until imaging.

**Table 1.** Information for primary antibodies.

| Primary Antibody | Manufacturer/Cat # | Dilution Factor |
| --- | --- | --- |
| Iba1 | Invitrogen, PA5-18039 | 1:1000 |
| Iba1 | Fujifilm Wako Pure Chemical Corporation,<br>019-19741 | 1:1000 |
| Clec7a (m-Dectin) | Invivogen, 6114-46-01 | 1:1000 |
| IL-4 | Invitrogen, 14-7108-81 | 1:500 |
| NeuN | Invitrogen, PA5-143567 | 1:1000 |
| TNFα | R&D system, MAB610 | 1:1000 |
| TOPRO | Biotium 40087 | 1:1000 |
| Methoxy X04 | Abcam, ab142818 | 1:75,000 |

**Table 2.** Information for secondary antibodies.

| Secondary Antibody | Manufacturer/Cat # | Dilution Factor |
| --- | --- | --- |
| anti-mouse IgG, Alexa Fluor 488 | Invitrogen, A21202 | 1:400 |
| anti-chicken IgG, Alexa Fluor 488 | Invitrogen, A78948 | 1:400 |
| anti-rabbit IgG, Alexa Fluor 488 | Invitrogen, A21206 | 1:400 |
| anti-rat IgG, Alexa Fluor 488 | Invitrogen, A21208 | 1:400 |
| anti-goat IgG, Alexa Fluor 546 | Life Technologies, A11056 | 1:400 |
| anti-rabbit IgG, Alexa Fluor 546 | Invitrogen, A10040 | 1:400 |
| anti-mouse IgG, Alexa Fluor 546 | Invitrogen, A10036 | 1:400 |
| anti-goat IgG, Alexa Fluor 546 | Invitrogen, A11056 | 1:400 |
| anti-chicken IgG, Alexa Fluor 647 | Invitrogen, A21449 | 1:400 |
| anti-rabbit IgG, Alexa Fluor 647 | Invitrogen, A31537 | 1:400 |
| anti-goat IgG, 647 | Abcam, ab150131 | 1:400 |
| anti-chicken 421 | JacksonImmuno Research, 703-6750155 | 1:200 |

### Microscopy and Image Analysis

Images were collected at The University of Chicago Integrated Light Microscopy core, on either a Olympus VS200 Slideview Research Slide Scanner using a 20x NA 0.8 Plan Apochromat objective (0.3251µm/pixel fluorescence), a Marianas Spinning Disk Confocal using a 40x NA 1.3 Plan Apochromat DIC and 63x / NA 1.0 W Plan Apochromat objective, or a Leica SP8 Laser Scanning Confocal using a 63x NA 1.4 HC Plan Apochromat objective. Z-stacks were collected with a 1-micron step-size (Slide Scanner, Marianas) or a 0.59-micron step size (SP8), and maximum intensity projections output using FIJI (NIH) before further analysis. All measures were quantified separately in cortex and hippocampus, unless otherwise noted using the freehand selection and threshold tools. Microglia coverage and activation at plaques, and internalized amyloid beta, were quantified using the “AND” function in the Image Calculator.

### Protein Isolation

Brain tissues were frozen and stored at –80°C until processing. Sequential extraction procedures were performed to isolate soluble and insoluble Aβ fractions.

### Soluble Aβ

Brains were first subset to hippocampus and cortex, and each were homogenized in RIPA buffer (Thermo Scientific, Cat 89900) supplemented with Halt^TM^ phosphatase and protease inhibitor cocktail (Thermo Scientific, Cat #7441) using a sonicator. Homogenates were kept on ice for 30 minutes, with vortexing every 3 minutes. The samples were then centrifuged at 13,200rpm for 30minutes at 4°C. The resulting supernatants were collected as soluble Aβ and stored in –80°C until further use.

### Insoluble Aβ

The pellet obtained from the soluble fraction extraction was resuspended in 70% formic acid containing phosphatase and protease inhibitor cocktail. The mixture was incubated for an hour at room temperature to solubilize insoluble proteins. The sample was then centrifuged at 16,000 rpm for 30 minutes at 4 °C, and the supernatant was collected as the insoluble Aβ fraction. Following extraction, the solution was neutralized with a 5M NaOH/1M Tris neutralization buffer. Proteins were precipitated using 9x volume of ice-cold 100% ethanol overnight, spun down in a centrifuge at 16,000 rpm for 30 minutes, and rinsed with 80% ethanol before spinning down again. The resulting pellet after precipitation was briefly air-dried and resuspended in RIPA buffer supplemented with phosphatase and protease inhibitor cocktail, collected as insoluble Aβ and stored at –80°C until further use.

### ELISA

Quantification of Aβ in both soluble and insoluble fractions was performed using a commercially available ELISA kit (Cell Signaling Technology, FastScan^TM^ β-Amyloid ELISA Kit #44261), following the manufacturer’s protocol. Homogenates were thawed on ice, and total protein concentration was determined using BCA protein assay kit (Thermo Scientific, Cat #23227). Based on the concentration, samples were diluted to a final concentration of 1ug for the soluble fraction and 1.5ug for insoluble fraction. Standards and diluted samples were loaded in duplicate into 96-well ELISA plates. After completion of the assay, absorbance was measured at 450nm using a microplate reader. Concentrations of Aβ were calculated using a standard curve generated from known concentrations of Aβ peptides provided by the manufacturer, averaged between technical replicates, and normalized to total protein content.

### Statistics

For microscopic analyses, all measures were first averaged within each section, then further averaged within each animal before analyses. Analyses were run using linear mixed effects modeling in the *lmer* package in R version 3.6.3, nested within animal for measures with multiple brain regions, and post-hoc analyses were run using *emmeans* in the *lmer* package. Multiple test corrections were done using Bonferroni (microglia coverage, cytokines, plaque occupancy, internalized amyloid, ELISA) or FDR (plaque burden and plaque coverage). For ELISA, resulting values were first normalized to µg per mg of protein so that insoluble and soluble fractions could be directly compared, then statistical analyses were performed similarly as above.

## Acknowledgements

We thank Dr. Sangram Sisodia for providing the initial APPPS1-21 mouse line for breeding, Benjamin Rutherford for assistance in mouse line maintenance, Carolina I. Castro-Rivera for helpful discussions on the source material, and the staff of the University of Chicago Light Microscopy Core for imaging assistance.

## Author Contributions

JVG and AJG designed the experiments. JVG and SWSW acquired and analyzed the data. JVG, SWSW and AJG interpreted the data. JVG drafted the manuscript. JVG, SWSW and AJG revised the manuscript. AJG oversaw the work.

## References

1. Ashley, E. A. (2016). Towards precision medicine. Nature Reviews Genetics, 17(9), 507–522. 10.1038/nrg.2016.86

2. Baligács, N., Albertini, G., Borrie, S. C., Serneels, L., Pridans, C., Balusu, S., & De Strooper, B. (2024). Homeostatic microglia initially seed and activated microglia later reshape amyloid plaques in Alzheimer’s Disease. Nature communications, 15(1), 10634. 10.1038/s41467-024-54779-w

3. Benzinger, T. L., Blazey, T., Jack, C. R., Koeppe, R. A., Su, Y., Xiong, C., Raichle, M. E., Snyder, A. Z., Ances, B. M., Bateman, R. J., Cairns, N. J., Fagan, A. M., Goate, A., Marcus, D. S., Aisen, P. S., Christensen, J. J., Ercole, L., Hornbeck, R. C., Farrar, A. M., Aldea, P., Barakos, J. A., Cruchaga, C., Davis, M., Reinwald, L. T., Liscic, R. M., Bracoud, L., Salloway, S., Schofield, P. R., Masters, C., Mayeux, R., Sperling, R., Johnson, K. A., Kaye, J., Quinn, J. F., Leverenz, J. B., Levey, A. I., Lah, J. J., Burns, D. K., Swerdlow, R. H., Brooks, W. S., Yesavage, J., Doody, R. S., Morris, J. C., & Dominantly Inherited Alzheimer Network. (2013). Regional variability of imaging biomarkers in autosomal dominant Alzheimer’s disease. Proceedings of the National Academy of Sciences, 110(45), E4502–E4509. 10.1073/pnas.1317918110

4. Bhatia, G., Ganesh, R., & Kulkarni, A. (2023). Cognitive impairment in opioid use disorders: Is there a case for use of nootropics?. Psychiatry research, 326, 115335. 10.1016/j.psychres.2023.115335

5. Billioti de Gage, S., Bégaud, B., Bazin, F., Verdoux, H., Dartigues, J. F., Pérès, K., Kurth, T., & Pariente, A. (2012). Benzodiazepine use and risk of dementia: Prospective population-based study. BMJ, 345, e6231. 10.1136/bmj.e6231

6. Bohannon, J. K., Hernandez, A., Enkhbaatar, P., Adams, W. L., & Sherwood, E. R. (2013). The immunobiology of toll-like receptor 4 agonists: from endotoxin tolerance to immunoadjuvants. Shock (Augusta, Ga.), 40(6), 451–462. 10.1097/SHK.0000000000000042

7. Calvo-Rodríguez, M., de la Fuente, C., García-Durillo, M., García-Rodríguez, C., Villalobos, C., & Núñez, L. (2017). Aging and amyloid β oligomers enhance TLR4 expression, LPS-induced Ca2+ responses, and neuron cell death in cultured rat hippocampal neurons. Journal of neuroinflammation, 14(1), 24. 10.1186/s12974-017-0802-0

8. Casali, B. T., MacPherson, K. P., Reed-Geaghan, E. G., & Landreth, G. E. (2020). Microglia depletion rapidly and reversibly alters amyloid pathology by modification of plaque compaction and morphologies. Neurobiology of disease, 142, 104956. 10.1016/j.nbd.2020.104956

9. Cui, Y., Liao, X. X., Liu, W., Guo, R. X., Wu, Z. Z., Zhao, C. M., Chen, P. X., & Feng, J. Q. (2008). A novel role of minocycline: attenuating morphine antinociceptive tolerance by inhibition of p38 MAPK in the activated spinal microglia. Brain, behavior, and immunity, 22(1), 114–123. 10.1016/j.bbi.2007.07.014

10. d’Errico, P., Ziegler-Waldkirch, S., Aires, V., Hoffmann, P., Mezö, C., Erny, D., Monasor, L. S., Liebscher, S., Ravi, V. M., Joseph, K., Schnell, O., Kierdorf, K., Staszewski, O., Tahirovic, S., Prinz, M., & Meyer-Luehmann, M. (2022). Microglia contribute to the propagation of Aβ into unaffected brain tissue. Nature neuroscience, 25(1), 20–25. 10.1038/s41593-021-00951-0

11. Do Carmo, S., & Cuello, A. C. (2013). Modeling Alzheimer’s disease in transgenic rats. Molecular neurodegeneration, 8, 37. 10.1186/1750-1326-8-37

12. Dublin, S., Walker, R. L., Jackson, M. L., Nelson, J. C., Weiss, N. S., Von Korff, M., & Jackson, L.A. (2011). Use of opioids or benzodiazepines and risk of pneumonia in older adults: a population-based case-control study. Journal of the American Geriatrics Society, 59(10), 1899–1907. 10.1111/j.1532-5415.2011.03586.x

13. Dufort, A., & Samaan, Z. (2021). Problematic Opioid Use Among Older Adults: Epidemiology, Adverse Outcomes and Treatment Considerations. Drugs & aging, 38(12), 1043–1053. 10.1007/s40266-021-00893-z

14. Eidson, L. N., & Murphy, A. Z. (2013). Blockade of Toll-like receptor 4 attenuates morphine tolerance and facilitates the pain relieving properties of morphine. The Journal of neuroscience: the official journal of the Society for Neuroscience, 33(40), 15952–15963. 10.1523/JNEUROSCI.1609-13.2013

15. Ferrini, F., Trang, T., Mattioli, T. A., Laffray, S., Del’Guidice, T., Lorenzo, L. E., Castonguay, A., Doyon, N., Zhang, W., Godin, A. G., Mohr, D., Beggs, S., Vandal, K., Beaulieu, J. M., Cahill, C. M., Salter, M. W., & De Koninck, Y. (2013). Morphine hyperalgesia gated through microglia-mediated disruption of neuronal Cl⁻ homeostasis. Nature neuroscience, 16(2), 183–192. 10.1038/nn.3295

16. Franchi, S., Moretti, S., Castelli, M., Lattuada, D., Scavullo, C., Panerai, A. E., & Sacerdote, P. (2012). Mu opioid receptor activation modulates Toll-like receptor 4 in murine macrophages. Brain, Behavior, and Immunity, 26(3), 480–488. 10.1016/j.bbi.2011.12.010

17. Fujii, K., Koshidaka, Y., Adachi, M., & Takao, K. (2019). Effects of chronic fentanyl administration on behavioral characteristics of mice. Neuropsychopharmacology reports, 39(1), 17–35. 10.1002/npr2.12040

18. Furcila, D., Domínguez-Álvaro, M., DeFelipe, J., & Alonso-Nanclares, L. (2019). Subregional Density of Neurons, Neurofibrillary Tangles and Amyloid Plaques in the Hippocampus of Patients With Alzheimer’s Disease. Frontiers in neuroanatomy, 13, 99. 10.3389/fnana.2019.00099

19. Gao, Y., Su, B., Ding, L., Qureshi, D., Hong, S., Wei, J., Zeng, C., Lei, G., & Xie, J. (2024). Association of Regular Opioid Use With Incident Dementia and Neuroimaging Markers of Brain Health in Chronic Pain Patients: Analysis of UK Biobank. The American journal of geriatric psychiatry: official journal of the American Association for Geriatric Psychiatry, 32(9), 1154–1165. 10.1016/j.jagp.2024.04.010

20. Gessi, S., Borea, P. A., Bencivenni, S., Fazzi, D., Varani, K., & Merighi, S. (2016). The activation of μ-opioid receptor potentiates LPS-induced NF-κB promoting an inflammatory phenotype in microglia. FEBS Letters, 590(17), 2813–2826. 10.1002/1873-3468.12313

21. Go, M., Kou, J., Lim, J. E., Yang, J., & Fukuchi, K. I. (2016). Microglial response to LPS increases in wild-type mice during aging but diminishes in an Alzheimer’s mouse model: Implication of TLR4 signaling in disease progression. Biochemical and biophysical research communications, 479(2), 331–337. 10.1016/j.bbrc.2016.09.073

22. Grace, P. M., Maier, S. F., & Watkins, L. R. (2015). Opioid-induced central immune signaling: implications for opioid analgesia. Headache, 55(4), 475–489. 10.1111/head.12552

23. Green, J. M., Sundman, M. H., & Chou, Y. H. (2022). Opioid-induced microglia reactivity modulates opioid reward, analgesia, and behavior. Neuroscience & Biobehavioral Reviews, 135, 104544. 10.1016/j.neubiorev.2022.104544

24. Griciuc, A., Patel, S., Federico, A. N., Choi, S. H., Innes, B. J., Oram, M. K., Cereghetti, G., McGinty, D., Anselmo, A., Sadreyev, R. I., Hickman, S. E., El Khoury, J., Colonna, M., & Tanzi, R. E. (2019). TREM2 acts downstream of CD33 in modulating microglial pathology in Alzheimer’s disease. Neuron, 103(5), 820–835.e7. 10.1016/j.neuron.2019.06.010

25. Hayashi, Y., Morinaga, S., Zhang, J., Satoh, Y., Meredith, A. L., Nakata, T., Wu, Z., Kohsaka, S., Inoue, K., & Nakanishi, H. (2016). BK channels in microglia are required for morphine-induced hyperalgesia. Nature communications, 7, 11697. 10.1038/ncomms11697

26. Hong, S., Beja-Glasser, V. F., Nfonoyim, B. M., Frouin, A., Li, S., Ramakrishnan, S., Merry, K. M., Shi, Q., Rosenthal, A., Barres, B. A., Lemere, C. A., Selkoe, D. J., & Stevens, B. (2016). Complement and microglia mediate early synapse loss in Alzheimer mouse models. Science, 352(6286), 712–716. 10.1126/science.aad8373

27. Hou, Y., Dan, X., Babbar, M., Wei, Y., Hasselbalch, S. G., Croteau, D. L., & Bohr, V. A. (2021). Ageing as a risk factor for neurodegenerative disease. Nature Reviews Neurology, 17(7), 392–406. 10.1038/s41582-019-0244-7

28. Hutchinson, M. R., Zhang, Y., Shridhar, M., Evans, J. H., Buchanan, M. M., Zhao, T. X., Slivka, P. F., Coats, B. D., Rezvani, N., Wieseler, J., Hughes, T. S., Landgraf, K. E., Chan, S., Fong, S., Phipps, S., Falke, J. J., Leinwand, L. A., Maier, S. F., Yin, H., Rice, K. C., … Watkins, L. R. (2010a). Evidence that opioids may have toll-like receptor 4 and MD-2 effects. Brain, behavior, and immunity, 24(1), 83–95. 10.1016/j.bbi.2009.08.004

29. Hutchinson, M. R., Lewis, S. S., Coats, B. D., Rezvani, N., Zhang, Y., Wieseler, J. L., Somogyi, A. A., Yin, H., Maier, S. F., Rice, K. C., & Watkins, L. R. (2010b). Possible involvement of toll-like receptor 4/myeloid differentiation factor-2 activity of opioid inactive isomers causes spinal proinflammation and related behavioral consequences. Neuroscience, 167(3), 880–893. 10.1016/j.neuroscience.2010.02.011

30. Hutchinson, M. R., Northcutt, A. L., Hiranita, T., Wang, X., Lewis, S. S., Thomas, J., van Steeg, K., Kopajtic, T. A., Loram, L. C., Sfregola, C., Galer, E., Miles, N. E., Bland, S. T., Amat, J., Rozeske, R. R., Maslanik, T., Chapman, T. R., Strand, K. A., Fleshner, M., Bachtell, R. K., … Watkins, L. R. (2012). Opioid activation of toll-like receptor 4 contributes to drug reinforcement. The Journal of neuroscience: the official journal of the Society for Neuroscience, 32(33), 11187–11200. 10.1523/JNEUROSCI.0684-12.2012

31. Islam, R., Choudhary, H. H., Zhang, F., Mehta, H., Yoshida, J., Thomas, A. J., & Hanafy, K. (2025). Microglial TLR4-Lyn kinase is a critical regulator of neuroinflammation, Aβ phagocytosis, neuronal damage, and cell survival in Alzheimer’s disease. Scientific reports, 15(1), 11368. 10.1038/s41598-025-96456-y

32. Kato, R. (1977). Drug metabolism under pathological and abnormal physiological states in animals and man. Xenobiotica, 7(1–2), 25–92. 10.3109/00498257709036242

33. Keren-Shaul, H., Spinrad, A., Weiner, A., Matcovitch-Natan, O., Dvir-Szternfeld, R., Ulland, T. K., David, E., Baruch, K., Lara-Astaiso, D., Tóth, B., Itzkovitz, S., Colonna, M., Schwartz, M., & Amit, I. (2017). A unique microglia type associated with restricting development of Alzheimer’s disease. Cell, 169(6), 1276–1290.e17. 10.1016/j.cell.2017.05.018

34. Kuo, Y. F., Westra, J., Harvey, E. P., & Raji, M. A. (2025). Use of Medications for Opioid Use Disorder in Older Adults. American journal of preventive medicine, 68(5), 1015–1021. 10.1016/j.amepre.2025.01.019

35. Lehnardt, S., Massillon, L., Follett, P., Jensen, F. E., Ratan, R., Rosenberg, P. A., Volpe, J. J., & Vartanian, T. (2003). Activation of innate immunity in the CNS triggers neurodegeneration through a Toll-like receptor 4-dependent pathway. Proceedings of the National Academy of Science (PNAS), 100:8514–8519. 10.1073/pnas.1432609100

36. Liang, X., Liu, R., Chen, C., Ji, F., & Li, T. (2016). Opioid system modulates the immune function: A review. Translational Perioperative and Pain Medicine, 1(1), 5–13.

37. Lin, T., Barash, J. A., Wang, S., Li, F., Yang, Z., Kofke, W. A., Sha, F., & Tang, J. (2025). Regular use of opioids and dementia, cognitive measures, and neuroimaging outcomes among UK Biobank participants with chronic non-cancer pain. Alzheimer’s & Dementia: The Journal of the Alzheimer’s Association, 21(5), e70177. 10.1002/alz.70177

38. Liu, X., Liu, B. L., Yang, Q., Zhou, X., & Tang, S. J. (2022). Microglial ablation does not affect opioid-induced hyperalgesia in rodents. Pain, 163(3), 508–517. 10.1097/j.pain.0000000000002376

39. Miao, J., Ma, H., Yang, Y., Liao, Y., Lin, C., Zheng, J., Yu, M., & Lan, J. (2023). Microglia in Alzheimer’s disease: Pathogenesis, mechanisms, and therapeutic potentials. Frontiers in Aging Neuroscience, 15, 1201982. 10.3389/fnagi.2023.1201982

40. Mika, J., Wawrzczak-Bargiela, A., Osikowicz, M., Makuch, W., & Przewlocka, B. (2009). Attenuation of morphine tolerance by minocycline and pentoxifylline in naive and neuropathic mice. Brain, behavior, and immunity, 23(1), 75–84. 10.1016/j.bbi.2008.07.005

41. O’Carroll, C., Fagan, A., Shanahan, F., & Carmody, R. J. (2014). Identification of a unique hybrid macrophage-polarization state following recovery from lipopolysaccharide tolerance. Journal of immunology (Baltimore, Md.: 1950), 192(1), 427–436. 10.4049/jimmunol.1301722

42. Pourhadi, N., Janbek, J., Gasse, C., Laursen, T. M., Waldemar, G., & Jensen-Dahm, C. (2024). Opioids and Dementia in the Danish Population. JAMA network open, 7(11), e2445904. 10.1001/jamanetworkopen.2024.45904

43. Prinz, M., Jung, S., & Priller, J. (2019). Microglia biology: One century of evolving concepts. Cell, 179(2), 292–311. 10.1016/j.cell.2019.08.053

44. Radde, R., Bolmont, T., Kaeser, S. A., Coomaraswamy, J., Lindau, D., Stoltze, L., Calhoun, M. E., Jäggi, F., Wolburg, H., Gengler, S., Haass, C., Ghetti, B., Czech, C., Hölscher, C., Mathews, P. M., & Jucker, M. (2006). Abeta42-driven cerebral amyloidosis in transgenic mice reveals early and robust pathology. EMBO reports, 7(9), 940–946. 10.1038/sj.embor.7400784

45. Rehm, J., Hasan, O. S. M., Black, S. E., Shield, K. D., & Schwarzinger, M. (2019). Alcohol use and dementia: a systematic scoping review. Alzheimer’s research & therapy, 11(1), 1. 10.1186/s13195-018-0453-0

46. Reed-Geaghan, E. G., Savage, J. C., Hise, A. G., & Landreth, G. E. (2009). CD14 and toll-like receptors 2 and 4 are required for fibrillar A{beta}-stimulated microglial activation. The Journal of neuroscience: the official journal of the Society for Neuroscience, 29(38), 11982–11992. 10.1523/JNEUROSCI.3158-09.2009

47. Reiss, D., Maduna, T., Maurin, H., Audouard, E., & Gaveriaux-Ruff, C. (2022). Mu opioid receptor in microglia contributes to morphine analgesic tolerance, hyperalgesia, and withdrawal in mice. Journal of Neuroscience Research, 100(1), 203–219. 10.1002/jnr.24626

48. Sasaguri, H., Nilsson, P., Hashimoto, S., Nagata, K., Saito, T., De Strooper, B., Hardy, J., Vassar, R., Winblad, B., & Saido, T. C. (2017). APP mouse models for Alzheimer’s disease preclinical studies. The EMBO Journal, 36(17), 2473–2487. 10.15252/embj.201797397

49. Shoff, C., Yang, T. C., & Shaw, B. A. (2021). Trends in Opioid Use Disorder Among Older Adults: Analyzing Medicare Data, 2013-2018. American journal of preventive medicine, 60(6), 850–855. 10.1016/j.amepre.2021.01.010

50. Shulman, M., Wai, J. M., & Nunes, E. V. (2019). Buprenorphine Treatment for Opioid Use Disorder: An Overview. CNS drugs, 33(6), 567–580. 10.1007/s40263-019-00637-z

51. Sobieraj DM, Baker WL, Martinez BK, et al. Comparative Effectiveness of Analgesics To Reduce Acute Pain in the Prehospital Setting [Internet]. Rockville (MD): Agency for Healthcare Research and Quality (US); 2019 Sep. (Comparative Effectiveness Review, No. 220.) Table 1, Onset, duration, and typical initial doses for analgesics, Available from: https://www.ncbi.nlm.nih.gov/books/NBK546195/table/ch2.tab1

52. Song, M., Jin, J., Lim, J. E., Kou, J., Pattanayak, A., Rehman, J. A., Kim, H. D., Tahara, K., Lalonde, R., & Fukuchi, K. (2011). TLR4 mutation reduces microglial activation, increases Aβ deposits and exacerbates cognitive deficits in a mouse model of Alzheimer’s disease. Journal of Neuroinflammation, 8, 92. 10.1186/1742-2094-8-92

53. Song, W. M., & Colonna, M. (2018). The identity and function of microglia in neurodegeneration. Nature Immunology, 19(10), 1048–1058. 10.1038/s41590-018-0212-1

54. Spedding, M., Jay, T., Costa e Silva, J., & Perret, L. (2005). A pathophysiological paradigm for the therapy of psychiatric disease. Nature reviews. Drug discovery, 4(6), 467–476. 10.1038/nrd1753

55. Subhramanyam, C. S., Wang, C., Hu, Q., & Dheen, S. T. (2019). Microglia-mediated neuroinflammation in neurodegenerative diseases. Seminars in Cell & Developmental Biology, 94, 112–120. 10.1016/j.semcdb.2019.05.004

56. Sun, N., Victor, M. B., Park, Y. P., Xiong, X., Scannail, A. N., Leary, N., Prosper, S., Viswanathan, S., Luna, X., Boix, C. A., James, B. T., Tanigawa, Y., Galani, K., Mathys, H., Jiang, X., Ng, A. P., Bennett, D. A., Tsai, L. H., & Kellis, M. (2023). Human microglial state dynamics in Alzheimer’s disease progression. Cell, 186(20), 4386–4403.e29. 10.1016/j.cell.2023.08.037

57. Upadhyay, J., Maleki, N., Potter, J., Elman, I., Rudrauf, D., Knudsen, J., Wallin, D., Pendse, G., McDonald, L., Griffin, M., Anderson, J., Nutile, L., Renshaw, P., Weiss, R., Becerra, L., & Borsook, D. (2010). Alterations in brain structure and functional connectivity in prescription opioid–dependent patients. Brain, 133(7), 2098–2114. 10.1093/brain/awq138

58. Vergadi, E., Vaporidi, K., & Tsatsanis, C. (2018). Regulation of Endotoxin Tolerance and Compensatory Anti-inflammatory Response Syndrome by Non-coding RNAs. Frontiers in immunology, 9, 2705. 10.3389/fimmu.2018.02705

59. Volkow, N. D., & Koob, G. F. (2015). Neurobiologic advances from the brain disease model of addiction. New England Journal of Medicine, 374(4), 363–371. 10.1056/NEJMra1511480

60. Warner, N. S., Hanson, A. C., Schulte, P. J., Habermann, E. B., Warner, D. O., & Mielke, M. M. (2022). Prescription opioids and longitudinal changes in cognitive function in older adults: A population-based observational study. Journal of the American Geriatrics Society, 70(12), 3526–3537. 10.1111/jgs.18030

61. Warner, N. S., Hanson, A. C., Schulte, P. J., Kara, F., Reid, R. I., Schwarz, C. G., Benarroch, E. E., Graff-Radford, J., Vemuri, P., Jack, C. R., Petersen, R. C., Warner, D. O., Mielke, M. M., & Kantarci, K. (2024). Prescription Opioids and Brain Structure in Community-Dwelling Older Adults. Mayo Clinic proceedings, 99(5), 716–726. 10.1016/j.mayocp.2024.01.018

62. Xie, N., Gomes, F. P., Deora, V., Gregory, K., Vithanage, T., Nassar, Z. D., Cabot, P. J., Sturgess, D., Shaw, P. N., & Parat, M. O. (2017). Activation of μ-opioid receptor and Toll-like receptor 4 by plasma from morphine-treated mice. Brain, behavior, and immunity, 61, 244–258. 10.1016/j.bbi.2016.12.002

63. Younger, J., McCue, R., & Mackey, S. (2009). Pain outcomes: a brief review of instruments and techniques. Current pain and headache reports, 13(1), 39–43. 10.1007/s11916-009-0009-x

64. Yu, J. T., Miao, D., Cui, W. Z., Ou, J. R., Tian, Y., Wu, Z. C., Zhang, W., & Tan, L. (2012). Common variants in toll-like receptor 4 confer susceptibility to Alzheimer’s disease in a Han Chinese population. Current Alzheimer research, 9(4), 458–466. 10.2174/156720512800492495

